# Reduction in neurons immunoreactive for parvalbumin, calretinin and calbindin in the anteroventral thalamic nuclei of individuals with Down syndrome

**DOI:** 10.1101/2024.02.01.578360

**Authors:** James C. Perry, Seralynne D. Vann

**Affiliations:** School of Psychology & Neuroscience and Mental Health Innovation Institute, Cardiff University, Cardiff, UK

**Keywords:** Anterior thalamic nuclei, amyloid, Alzheimer’s disease, calcium-binding proteins, immunohistochemistry

## Abstract

The anterior thalamic nuclei are important for cognition, and memory in particular. However, little is known about how the anterior thalamic nuclei are affected in many neurological disorders partly due to difficulties in selective segmentation in *in vivo* scans, due to their size and location. Post-mortem studies, therefore, remain a valuable source of information about the status of the anterior thalamic nuclei. We used post-mortem tissue to assess the status of the anteroventral thalamic nucleus in Down syndrome using samples from males and females ranging from 22-65 years in age and comparing to tissue from age matched controls. As expected, there was increased beta-amyloid plaque expression in the Down syndrome group. While there was a significant increase in neuronal density in the Down syndrome group, the values showed more variation consistent with a heterogeneous population. The surface area of the anteroventral thalamic nucleus was smaller in the Down syndrome group suggesting the increased neuronal density was due to greater neuronal packing but likely fewer overall neurons. There was a marked reduction in the proportion of neurons immunoreactive for the calcium-binding proteins calbindin, calretinin, and parvalbumin in individuals with Down syndrome across all ages. These findings highlight the vulnerability of calcium-binding proteins in the anteroventral nucleus in Down syndrome, which could both be driven by, and exacerbate, Alzheimer-related pathology in this region.

## Introduction

Down syndrome is the most common chromosomal disorder, affecting 1 in 1000 live births in the UK. This highlights the need to have a better understanding of how the brain can be affected with the long-term goal of better supporting cognitive function and reducing age-related decline. Memory is often affected in individuals with Down syndrome and this has typically been attributed to structural changes in the hippocampal formation (Carducci et al., 2013, Kemper 1991, Pinter et al., 2001). However, other brain regions that are important for memory, including the anterior thalamic nuclei, also appear to be compromised in Down syndrome; for example, we previously found a marked reduction of neurons in the anterior thalamic nuclei in individuals with Down syndrome (Perry et al., 2019). While these findings implicate the anterior thalamic nuclei in Down syndrome, a limitation of that study was that the tissue was exclusively from older females, so it is unclear whether these changes reflect the presence of advanced Alzheimer pathology in this group, rather than being more specifically related to Down syndrome. Alzheimer-related pathology is a common feature in individuals with Down syndrome with the onset of amyloid pathology commencing as early as the late teens and dementia often occuring by the mid 50s (Davidson et al., 2018, Head et al., 2012).

One aim of the current study was to determine whether the anterior thalamic nuclei exhibit changes in Down syndrome at younger ages. We, therefore, analysed post-mortem tissue from 15 individuals with Down syndrome (6 female) ranging from 22 to 62 years and compared this to tissue from 12 control individuals (6 female) ranging in age from 38 to 70 years. We measured cell density in the anteroventral thalamic nucleus, including neurons and glial cells, as well as the numbers of cells that stained positive for the calcium-binding proteins, calretinin, parvalbumin and calbindin, to determine whether any of these cell populations appeared to be affected in individuals with Down syndrome. The number of neurons immunopositive for calcium-binding proteins is generally reduced in Alzheimer’s disease (Satoh et al., 1991). A reduction in calbindin and parvalbumin stained neurons has also been reported in the cortex of older participants with Down syndrome (Kobayashi et al., 1990) but there is relatively little known about the expression of calcium-binding proteins in the human anterior thalamic nuclei in general, and even less knowledge in the case of Down syndrome. Calcium-binding proteins are important for calcium homeostasis and intracellular signalling (Fairless et al., 2019), highlighting the importance of understanding their expression in Down syndrome.

A subsequent aim was to assess whether Alzheimer’s disease related pathology contributed to neuronal calcium-binding protein expression given the links between calcium homeostasis and both beta-amyloid (Aβ) and tau (Guan et al., 2021). Diffuse Aβ plaques are comprised primarily of Aβ42 and develop at a younger age, with the earliest cortical deposits being found in Down syndrome around age 12 (Lemere et al., 1996). In contrast, the more mature plaques containing Aβ40 (Iwatsubo et al., 1995), and neurofibrillary tangles, have been more typically reported in cortical and parahippocampal regions from age 30-35 in individuals with Down syndrome (Lemere et al., 1996, Head et al., 2003). To determine how the differential development of plaques and tangles relate to expression of calcium-binding proteins, we processed the tissue for tau, Aβ42, Aβ40, and Aβ-4G8 to visualize total amyloid burden (Mandler et al., 2014). A more comprehensive understanding of the vulnerability of cells within the anteroventral thalamic nucleus will help identify possible contributing factors underlying structural changes and the extent to which these are driven by Alzheimer-related pathology.

## Methods

### Tissue samples

Samples originated from formalin fixed paraffin embedded tissue blocks from the brains of 15 people with Down syndrome (age range 22 – 65 years) and 12 controls (age range 38 – 70). Slide-mounted consecutive serial coronal sections cut at 6µm thickness were provided by the UK Brain Banks Network (UKBBN). The UKBBN adheres to agreed common standards, therefore, handling of tissue was comparable across sites. Details of each case are provided in Additional File 1: Supplementary Table 1. This study was conducted in accordance with the UK Human Tissue Act (2004). Approval for the study was given by the School of Psychology Research Ethics Committee, Cardiff University, and use of the tissue was covered by the generic approval from each brain bank (Manchester, REC reference 19/NE/0242; London, 23/WA/0124; Cambridge, 20/EE/0283; Oxford, 23/SC/0241).

**Table 1.** Primary antibodies used for immunohistochemistry.

| Antigen | Antibody | Dilution | Detection | Source |
| --- | --- | --- | --- | --- |
| Aβ, 17-24 | Mouse anti-4G8 (monoclonal) | 1:1000 | Streptavidin/DAB | BioLegend 800708 |
| Aβ, 1-40 | Rabbit anti- β1-40 (polyclonal) | 1:200 | ImmPress polymer horse-anti-rabbit HRP/StayYellow | Merck, AB5074P |
| Aβ, 1-42 | Mouse anti-124F (monoclonal) | 1:500 | ImmPress polymer horse-anti-mouse AP/StayBlue | BioLegend, 805509 |
| Tau (pSer202/ PThr205) | Rabbit anti-AT8/AH36 (monoclonal) | 1:500 | Streptavidin/DAB | StressMarq Biosciences, SMC-601. |
| Parvalbumin | Recombinant rabbit anti-parvalbumin (monoclonal) | 1:500 | Streptavidin/DAB | Abcam, ab181086 |
| Calretinin | Goat anti-calretinin (polyclonal) | 1:100 | ImmPress polymer horse-anti-goat HRP/DAB | Novus, AF5065 |
| Calbindin | Chicken anti-calbindin D-18K (polyclonal) | 1:250 | Streptavidin/DAB | Novus, NBP2-50028 |

### Immunohistochemistry

For each immunohistochemistry procedure, all sections were processed as a single batch with staining carried out within a single session. Sections were deparaffinized in xylene then rehydrated in a descending series of alcohols. Endogenous enzyme activity was quenched in either methanol containing 3% H_2_O_2_ or Bloxall solution (SP-6000-100, Vector Laboratories, US) to block both peroxidase and alkaline phosphate activity. Epitope retrieval was performed by incubating sections in 98% formic acid for 5 mins. Sections were then washed three times in 0.1M PBS containing 0.2% triton-X 100 followed by blocking in 10% normal goat or horse serum (Vector Laboratories, US) for 1 h at room temperature. Sections were incubated in primary antibody solution overnight at 4°C in a humidified chamber (Table 1). The Aβ-4G8, AT8, parvalbumin, and calbindin samples were incubated in an appropriate biotinylated secondary antibody solution (1:200 dilution) for 1 h at room temperature followed by streptavidin horseradish peroxidase solution for 30 mins. The calretinin samples were incubated in ImmPRESS horseradish peroxidase polymer detection kit for 30mins (Vector Laboratories, US). Samples were developed in a 3,3’-diaminobenzidine (DAB) substrate kit (Vector Laboratories, US), counterstained using QS haematoxylin (Vector Laboratories, US), then dehydrated in an ascending series of alcohols, cleared in xylene and coverslipped using DPX mounting media. The Aβ 42/40 double-labelled samples were incubated in ImmPRESS horseradish peroxidase polymer then ImmPRESS alkaline phosphate polymer kits for 30 mins each (Vector Laboratories, US). For these sections, Aβ40 was visualized using StayYellow HRP chromogen (ab169561, Abcam,UK) followed by Aβ42 visualization using StayBlue AP chromogen (ab176915, Abcam, UK), counterstained with haematoxylin, air dried and coverslipped using aqueous mounting media. For each case, one section was processed for each immunohistochemical marker except for one 70-year-old control case where tissue was not stained for calbindin and Tau AT8 due to insufficient samples. Primary antibody specificity was assessed by omitting the primary from one sample per marker. In addition, the expected regional specificity of the neuropil stain for calretinin, parvalbumin, and calbindin at low magnification along with neuronal, but not glial cell, label at higher magnification was confirmed.

### Imaging and analysis

All analyses of the anterior thalamus in this study employed digital pathology methods. This allowed objective assessments of the whole region on each section. To do this, images of the anterior thalamus were obtained using an automated slide scanner (Olympus VS200, Olympus Life Science, Japan) equipped with a color iDS camera (Olympus VS-264C). The anterior thalamus was imaged using a 40x objective (NA 0.95) for Aβ-4G8, Tau AT8, hematoxylin, calretinin, parvalbumin and calbindin and a 20x objective (NA 0.8) for Aβ 40/42. The images were loaded into QuPath (v0.4.3, Bankhead et al., 2017) and the anterior ventral thalamic nuclei was annotated as previously described (Perry et al., 2019). The percentage area of the anteroventral thalamic nucleus occupied by Aβ-4G8, Aβ 40/42, and Tau AT8 was calculated by training a pixel classification model to identify the relative area occupied by each marker.

For neurons and glia cells, the StarDist deep-learning model (Schmidt et al., 2018) was used to identify each stained object in sections stained with hematoxylin alone. An object classifier was trained to identify each object as either a neuron, astrocyte, oligodendrocyte, or microglia. The classifier employed several measures including area, length, circularity, diameter, hematoxylin intensity and a 100µm weighted smoothing for each measure. Similarly, StarDist was used to detect calretinin, parvalbumin, and calbindin positive neurons in DAB and hematoxylin counter-stained sections. To do this, the DAB and hematoxylin channels were extracted and summed into a single detection channel. Positive neurons were identified by filtering detections to a minimum nucleus area, circularity, DAB and hematoxylin intensity. To provide an indication of the performance of each cell classifier, one expert manually annotated a 1,000,000 µm^2^ area on one section for hematoxylin stained neurons and glia and a 4,000,000µm^2^ area on one section per calcium binding protein marker for samples from both the controls and people with Down syndrome. The manually annotated sections were regarded as the ground truth and were used to compare the accuracy, precision, and recall of each cell classifier. Each cell classifier performed well, with high accuracy (range: 89-99%), precision (range 89-98%), and recall (range: 89-97%) achieved, with comparable performance for samples from both the control and Down syndrome groups (see Additional File 1: Supplementary Table 2).

**Table 2.**
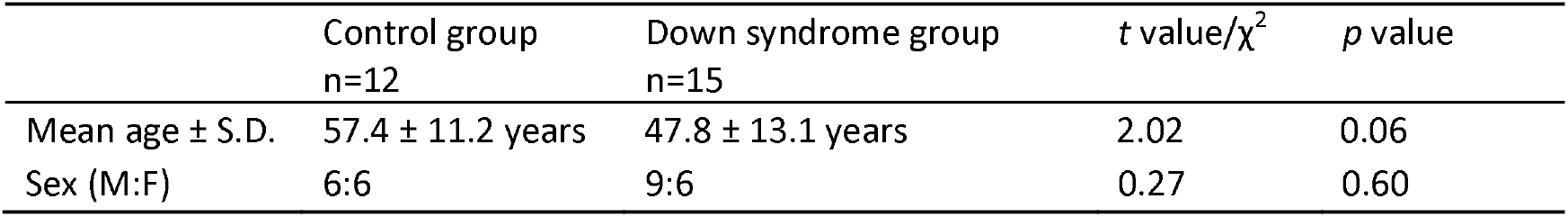
Demographic data.

| | Control group<br>n=12 | Down syndrome group<br>n=15 | *t* value/$\chi^2$ | *p* value |
| --- | --- | --- | --- | --- |
| Mean age ± S.D. | 57.4 ± 11.2 years | 47.8 ± 13.1 years | 2.02 | 0.06 |
| Sex (M:F) | 6:6 | 9:6 | 0.27 | 0.60 |

Neuron and glia counts could not be obtained from DAB-stained sections that were counterstained with hematoxylin due to the background staining from DAB. As such, neuron and glia measurements were only acquired from the hemaotxylin-only stained sections (one section per case). The numbers of microglia were added to the total glia density and not analysed alone due to low numbers. Calbindin, calretinin and parvalbumin counts were all derived from separate brain DAB-stained sections, and again from one section per case. For neuron and glial counts, a measure of the total number of items was derived within the boundaries of the anteroventral thalamic nuclei on each section, these measures were then adjusted for the area of the anteroventral thalamic nucleus on the section, providing a measure of objects per mm^2^. Calbindin, calretinin and parvalbumin counts were further adjusted for neuronal density for each case, to provide an approximate proportion of immunopositive neurons.

One 40x annotated image, selected from the middle of the available consecutive series of sections (i.e., the largest available region for each case) was used to provide an approximate estimate of the cross-sectional area of the anteroventral thalamic nucleus (AV). Each section was coded as either anterior or posterior AV, as the anteroposterior levels varied across samples.

### Data Analysis

Statistical analyses were performed in JASP version 0.18.2 (JASP Team, 2024). Independent means *t*-tests were used to compare the age and post-mortem delay of controls and individuals with Down syndrome. The distribution of gender across the groups was assessed using a Chi-squared test. Pearson correlation was used to assess whether post-mortem delay, i.e. the time between death and tissue fixation, independent of group, was correlated with any measure. A 2 x 2 factorial ANOVA was used to assess the cross-sectional area of the AV with Diagnosis (Control vs Down syndrome) as the between groups factor and the Anterior-posterior level (anterior vs posterior) of the AV as the within groups factor. Comparisons between histological markers across groups were carried out using independent means *t*-tests. One-tailed tests were used for Aβ pathology markers while two-tailed test were used for all other measures. Welch’s t-test for unequal variances was used wherever appropriate. A Mann-Whitney *U* test was used when normality was not assumed. Wherever relevant, effect sizes between groups were expressed as Hedges *g* when equal variances were assumed, Glass’s Δ when equal variances were not assumed, rank-biserial correlation following a Mann-Whitney *U* test, or 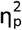 following factorial ANOVA. Separate multiple regressions with backward elimination (*p* <0.1 retention) were modelled with neuronal density, calretinin, parvalbumin, and calbindin proportions as the dependent variables, and Aβ42, diagnosis, and age as the co-variates. Among the Aβ pathology markers stained in this study, Aβ42 was selected as a predictor for the regression model as it is thought to be the more pathogenic form of Aβ (Bodani et al., 2015, Iwatsubo et al., 1995). Age was included in the model due to the association between advancing age and AD pathology in Down syndrome (Davidson et al., 2018). Tau was not included due low expression. The most parsimonious model with the best fit is reported. Additional details of regression summaries, ANOVAs, and coefficients for each model are included in Additional File 2: Supplementary tables 3 -14. Multiple comparisons were corrected with the Benjamini-Hochberg procedure (Q = 0.05). The corrected *p* values (*p*_*c*_) of less than 0.05 were considered significant. Given the small numbers, correlations with age were not carried out but the data are presented in scatterplots so that age and gender distribution across groups can be visualized.

In addition to significance testing, the Bayesian *t*-test framework was applied to the key outcome measures: proportion of calretinin, parvalbumin, and calbindin positive neurons. Bayesian Mann-Whitney *U* test, with Markov Monte Carlo set to five chains of 10, 000 iterations, was used when normality was not assumed. The standardised effect size (δ) was assigned a Cauchy prior distribution with *r* = 1/√2. For each test, the Bayes factor (BF_10_), was calculated to quantify the relative evidence for the null hypothesis (H_0_) compared to the alternative hypothesis (H_1_). BF_10_ provides a continuous measure of evidence where a BF_10_ between 1 and 3 is considered anecdotal evidence, between 3 and 10 is considered moderate evidence and BF_10_ greater than 10 is considered strong evidence, in favour of the alternative hypothesis, respectively. In comparison, a BF_10_ between 1 and 0.33 is considered anecdotal evidence, between 0.33 and 0.1 is considered moderate evidence and lower than 0.1 can be interpreted as strong evidence, in favour of the null hypothesis, respectively (van Doorn et al., 2021). Here it is important to note that verbal labels used to categorise different Bayes factors can be useful to facilitate scientific communication, but caution is needed due to the arbitrary nature of these labels and the continuous nature of the Bayes factor. The Bayesian t test and Bayesian Mann-Whitney *U* test posterior and prior plots, along with robustness checks where applicable are included in Additional File 3: Supplementary figures 1 -3. Finally, we applied the Bayesian framework to the calcium binding protein regression analyses as described above. In addition to providing the BF_10_ for the most parsimonious model, this framework also quantifies the relative evidence for the inclusion (BF_inclusion_) of each co-variate in the model. A beta prior probability (a = b =1) was applied to the models and a JZS prior (r scale = 0.354) was applied to the regression coefficients. Additional details of Bayesian regression model comparisons, posterior summaries of coefficients, and descriptive statistics are included in Additional File 4: Supplementary tables 15 – 23. Figures were produced in R using ggplot2 (Wickham, 2016).

**Fig 1.**
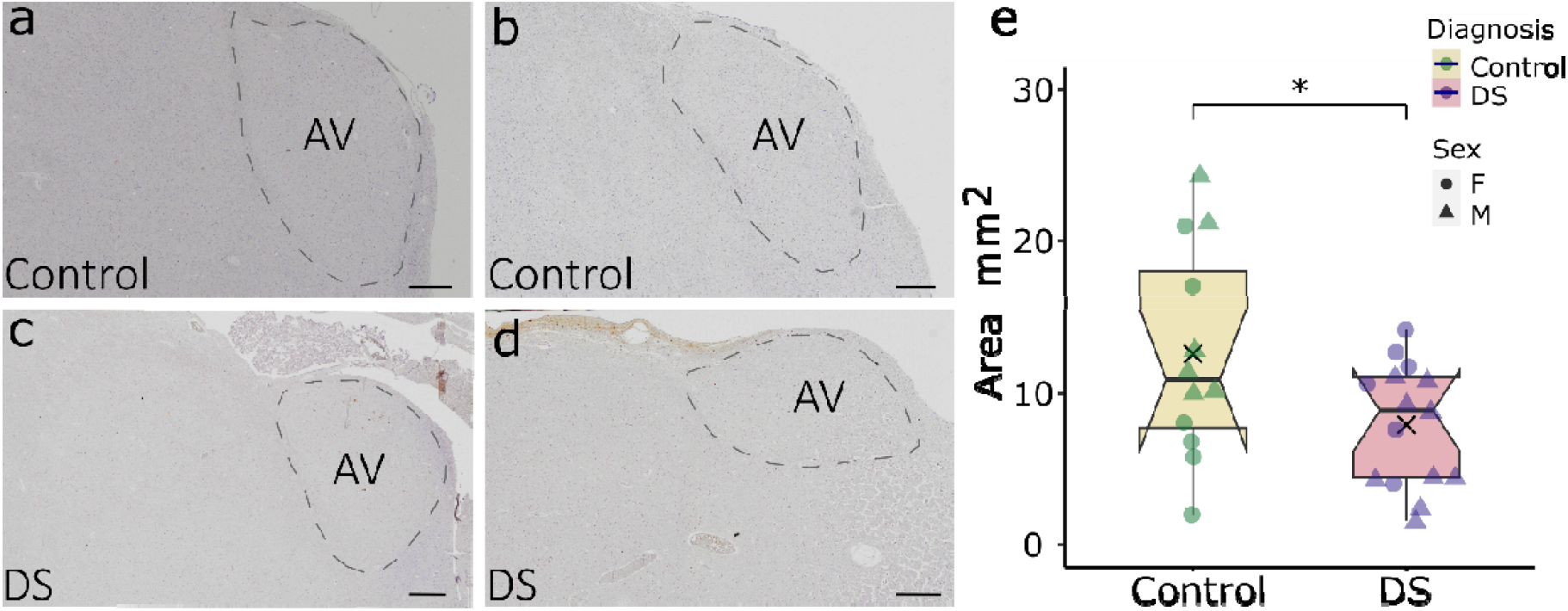
Cross-sectional area of the anteroventral thalamic nucleus. Representative examples of the anteroventral thalamic nucleus from similar middle anterior – posterior levels in (a) 63yo male control (case# BBN110.35427), (b) 61yo female control (case# BBN110.28899), (c) 47yo male with Down syndrome (case# BBN_17189), (d) 60yo female with Down syndrome (case# BBN_2990) and (e) box plot of the cross-sectional area of the anteroventral thalamic nuclei. The box indicates the inter-quartile range (IQR), the whiskers show the range of values that are within 1.5 x IQR, the horizontal line indicates the median, and the cross indicates the mean. The notches represent the 95% confidence interval for each median (1.58 x IQR/√(n)). Control, control group; DS, Down syndrome group, F, females, M, males; *p<0.05, **p<0.01, ***p<0.001. Scale bar = 1mm. Note: The examples of the anteroventral thalamic nucleus in (a) to (d) are all oriented with dorsal towards the top of the image. The orientation of the anteroventral nucleus can vary between individuals at similar anterior – posterior levels.

## Results

### Group demographics

The control group tissue came from a slightly older population (range 38-70 years) than the Down syndrome (DS) group tissue (range 22-62 years), although the difference did not reach significance (Table 2). There was no difference in gender distribution across the two groups (see Table 2).

Given the sample size and distribution of data, we did not look at specific correlations between histological measures and age. While there were no effects of gender, due to the small numbers it is possible that more subtle gender effects may not be detected, therefore, age and gender distributions are presented graphically for the histological measures. Post-mortem delay data were available for all 12 controls and 7 of the 15 Down syndrome samples. While there was a trend for individuals with Down syndrome (M = 70.7h, SD = 28.6) to have a longer post-mortem delay than controls (M = 54.2h, SD = 21.7), this difference was not significant t(17) = -1.43, *p* = 0.17, uncorrected, g = 0.47, 95% CI: [-1.60, 0.32]. Pearson correlation was used to assess whether the post-mortem delay, independent of group, was correlated with each measure. While there was no significant correlation between post-mortem delay and any measure, there was a moderate relationship between post-mortem delay and calretinin density (*r* = -0.45, *p* = 0.06, uncorrected; see Additional File 1: Supplementary Table 3).

### Regional area of anteroventral thalamic nucleus

While we did not have the tissue necessary to derive volumetric measurements for the anteroventral thalamic nucleus (AV), we did estimate the cross-sectional regional area from one mid-section per case (Fig. 1a - d). We found that the area of AV was approximately 37% smaller in the DS group compared to controls and this difference was significant as shown by a main effect of Diagnosis 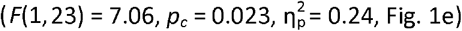. The smaller AV area in the DS group was not due to differences in the anterior-posterior levels between the samples as neither the main effect of Anterior-posterior level 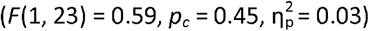 nor the Diagnosis x Anterior-posterior level interaction were significant 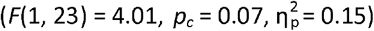.

### Neurons and glia

There was increased neuron density in the DS group (*t*(16.482) = -2.27, p_c_ = 0.049, Δ = -0.61, 95% CI: [-1.39, 0.19], Fig. 2a, e, i), however, the range in neuron density was much greater in the DS group (44-288 cells/mm^2^ ) than in the control group (74-129 cells/mm) suggesting a more heterogeneous population. While the density of total glia was slightly higher in the DS group, this difference was not significant (t(25) = -1.13, p_c_ = 0.30, g = -0.423, 95% CI: [-1.19, 0.35], Fig. 2b, f, j). The numerical increase in total glia density was driven by a significant increase in astroglia density (*t*(25) = -2.33, *p*_*c*_ = 0.04, *g* = -0.88, 95% CI: [-1.67, -0.07], Fig. 2c, g, k), but not, oligodendrocyte density (*t*(25) = 0.81, *p*_*c*_ = 0.45, *g* = 0.30, 95% CI: [-0.46, 1.06], Fig. 2d, h, l), which was comparable across the groups.

**Fig 2.**
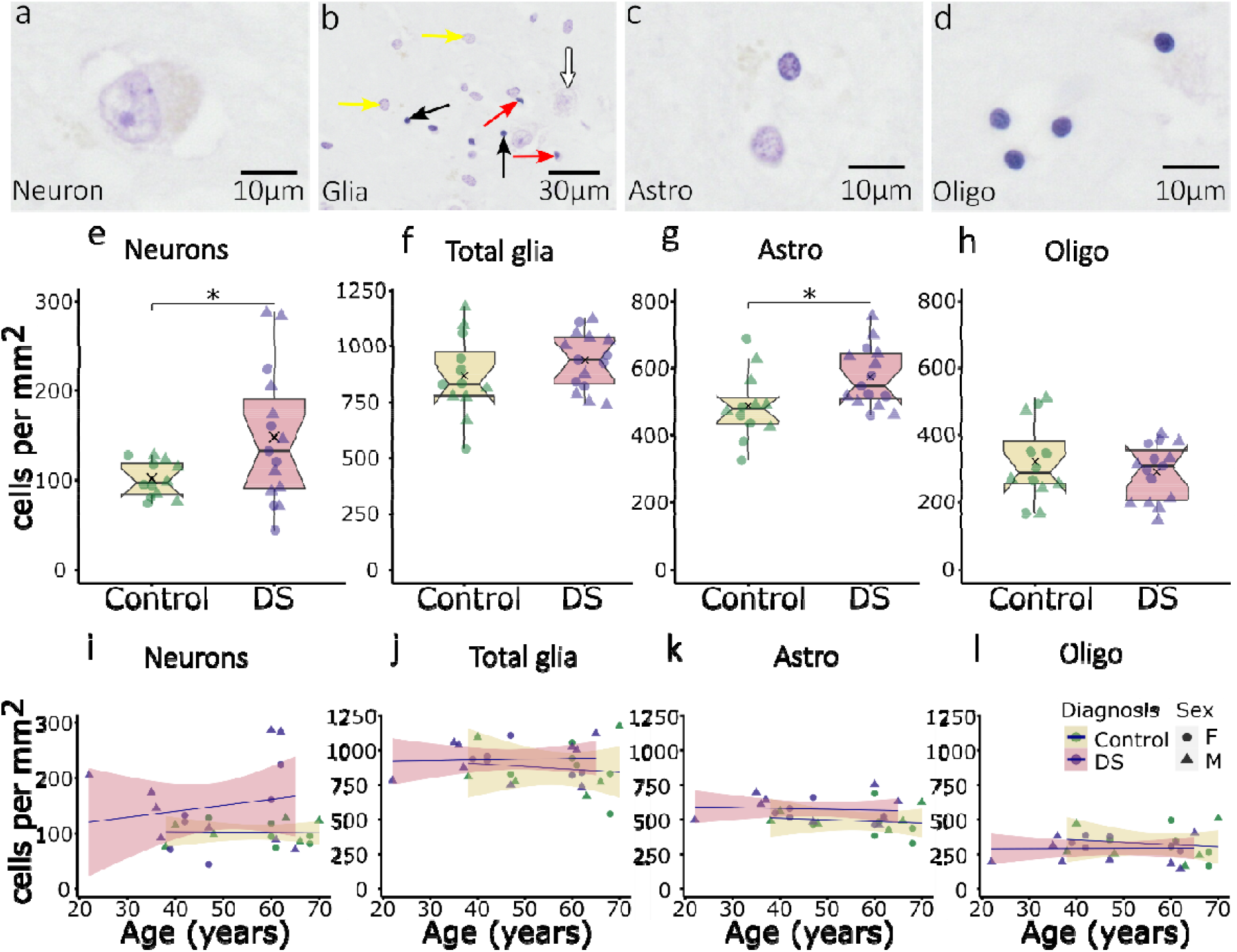
Neuron and glia density in the anteroventral thalamic nucleus. Top panel, hematoxylin stained examples of (a) a neuron with clearly stained nucleolus, nucleus, and surrounded by cytoplasm, (b) examples of astroglia (yellow arrows), oligodendrocytes (black arrows), comma shaped microglia (red arrow), and neurons (white arrow) and zoomed in examples of (c) astroglia showing a typical pale nucleus with heterochromatin staining concentrated in granules below the nuclear membrane and (d) oligodendrocytes with a classic rounded nucleus and dense chromatin staining. Middle panel, box plots of the densities of (e) neurons, (f) total glia, (g) astroglia and (h) oligodendrocytes. The box indicates the inter-quartile range (IQR), the whiskers show the range of values that are within 1.5 x IQR, the horizontal line indicates the median, and the cross indicates the mean. The notches represent the 95% confidence interval for each median (1.58xIQR/√(n)). Bottom panel, scatterplot showing the relationship between age and density of (i) neurons, (j) total glia, (k) astroglia and (l) oligodendrocytes. The lines are fitted with the general linear model and the shaded areas are the 95% confidence interval. Control, control group; DS, Down syndrome group, F, females, M, males. *p<0.05, **p<0.01, ***p<0.001. Note: All neuron and glia examples come from the same 61yo male control (case# BBN110.28899). Samples were imaged at 40x magnification and zoomed in using QuPath to show the features of each cell type.

### Amyloid and tau

As expected, the area fraction of AV occupied by Aβ was significantly higher in the DS group than the control group for all three markers: Aβ-4G8 (Fig. 3a, d), Aβ42 (Fig. 3b, e) and Aβ40 (Fig. 3b, e) [Aβ-4G8: Mann-Whitney *U* = 15.00, *p*_*c*_ < 0.001, *r* = -0.83, 95% CI: [-∞, -0.68], Fig. 3g, i; Aβ42: *t*(14.03) = - 4.32, *p*_*c*_ = 0.002, Δ = -1.12, 95% CI: [-∞, -0.38], Fig. 3h, j; Aβ40: Mann-Whitney *U* = 49.50, *p*_*c*_ = 0.02, *r* = -0.45, 95% CI: [-∞, -0.11]]. While there was a significant group difference for Aβ40, expression was generally low with levels near zero for 13 of the 15 tissue samples from the DS group. However, moderate expression was found in one case (47-year-old female, 1.14% coverage), and high expression was found in one case (47-year-old male, 7.24% coverage). Tau AT8 expression in the AV was generally low and did not differ between the groups (*p* = 0.36, uncorrected). Tau AT8 expression was near zero in 12 of the 15 tissue samples from individuals with DS. However, moderate expression was found in one case (65-year-old male: 1.7% coverage), and high expression was found in two cases (47-year-old female, 9.1% coverage; 60-year-old male, 6.5% coverage; Fig. 3c, f).

**Fig 3.**
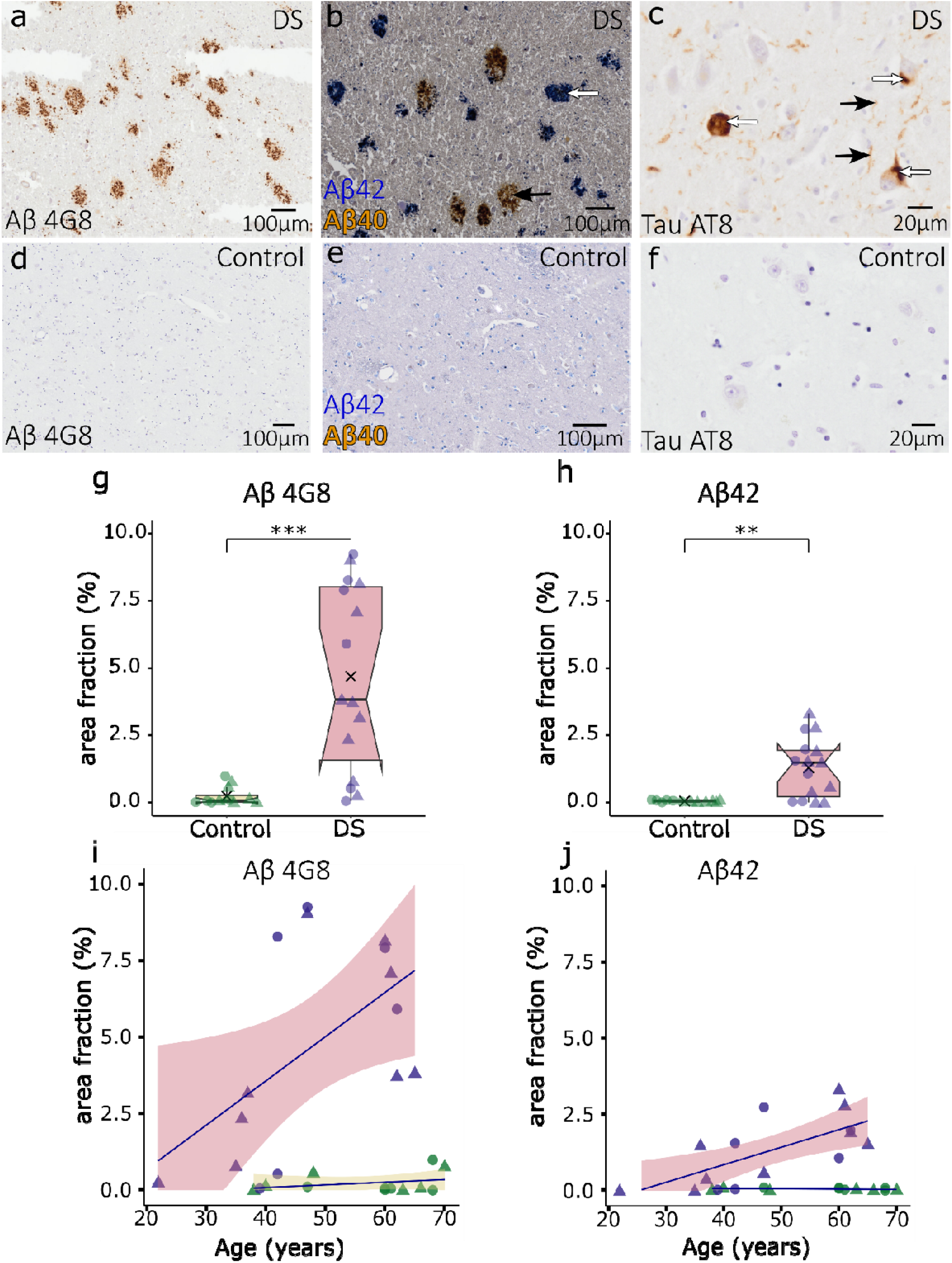
Alzheimer’s disease pathology in the anteroventral thalamic nucleus. Top panel, examples of (a) Aβ-4G8 plaques, (b) blue stained Aβ42 (white arrow) and yellow stained Aβ40 (black arrow) plaques taken from the same area of the anteroventral thalamic nucleus from a 47yo female with Down syndrome (case# BBN_17186) and (c) Tau AT8 positive neurofibrillary tangles (white arrows) and neuropil threads (black arrows) taken from a 65yo male with Down syndrome (case# BBN_2975). Second panel, no expression of (d) Aβ-4G8, (e) Aβ42 and Aβ40, and (f) Tau AT8 in a representative control taken from a 61yo female control (case# BBN110.28899). Third panel, box plots of the percent area fraction occupied by (g) Aβ 4G8 and (h) Aβ42. The box indicates the inter-quartile range (IQR), the whiskers show the range of values that are within 1.5xIQR, the horizontal line indicates the median, and the cross indicates the mean. The notches represent the 95% confidence interval for each median (1.58xIQR/√(n)). Bottom panel, scatterplot showing the relationship between age and the percent area fraction occupied by (i) Aβ 4G8 and (j) Aβ42. The lines are fitted with the general linear model and the shaded areas are the 95% confidence interval, truncated at zero. Control, control group, DS, Down syndrome group; F, females, M, males. *p<0.05, **p<0.01, ***p<0.001.

### Calcium-binding proteins (calretinin, calbindin, parvalbumin)

Counts of immunopositive neurons for each of the calcium-binding proteins were normalized by dividing each calcium-binding protein marker density by the total neuronal density for each individual case, to provide a relative proportion of immunopositive neurons. The proportion of neurons immunopositive for all three calcium-binding proteins in AV was significantly reduced in the DS group [calretinin, *t*(25) = 3.23, *p*_*c*_ = 0.008, *g* = 1.21, 95% CI: [0.37, 2.03], Fig. 4a, d, g, j; parvalbumin, Mann-Whitney *U* = 149, *p*_*c*_ = 0.009, *r* = 0.66, 95% CI: [0.33, 0.84], Fig. 4b, e, h, k; calbindin, Mann-Whitney *U* = 156, *p*_c_ < 0.001, *r* = 0.89, 95% CI: [0.75, 0.96], Fig. 4c, f, i, l]. The same pattern of results was found when using the density of neurons immunopositive for calcium-binding proteins in the AV, rather than proportion [calretinin, *t*(25) = 2.61, *p*_*c*_ = 0.023, *g* = 0.98, 95% CI: [0.16, 1.78]; parvalbumin, Mann-Whitney *U* = 144, *p*_*c*_ = 0.018, *r* = 0.60, 95% CI: [0.24, 0.81]; calbindin, Mann-Whitney *U* = 140, *p*_*c*_ = 0.006, *r* = 0.70, 95% CI: [0.38, 0.87]].

**Fig 4.**
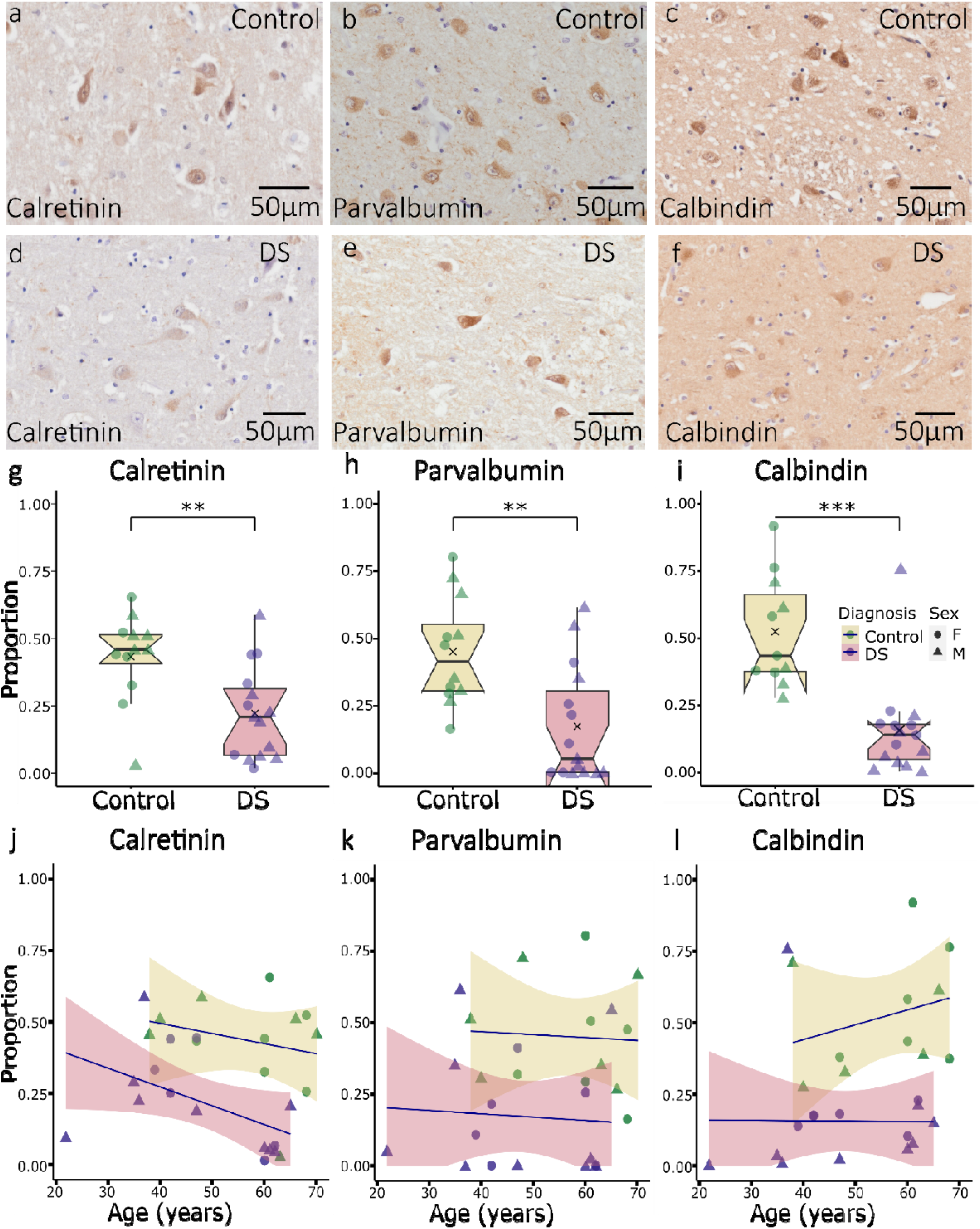
Calcium-binding proteins in the anteroventral thalamic nucleus. Top panel, representative examples of (a) calretinin from a 66yo female control (case# BBN_24381), (b) parvalbumin from a 60yo female control (case# BBN110.28668), and (c) calbindin positive neurons from a 61yo female control (case# BBN110.28899). Second panel, representative examples of (d) calretinin from a 37yo male with DS (case# BBN002.28703), (e) parvalbumin from a 60yo female with DS (case# BBN_2990), and (f) calbindin from a 37yo male with DS (case# BBN002.28703). Third panel, box plots of normalized densities of (g) calretinin, (h) parvalbumin, and (i) calbindin positive neurons. The box indicates the inter-quartile range (IQR), the whiskers show the range of values that are within 1.5xIQR, the horizontal line indicates the median, and the cross indicates the mean. The notches represent the 95% confidence interval for each median (1.58xIQR/√(n)). Bottom panel, scatterplot showing the relationship between age and normalized densities of (j) calretinin, (k) parvalbumin, and (l) calbindin positive neurons. The lines are fitted with the general linear model and the shaded areas are the 95% confidence interval, truncated at zero. Control, control group; DS, Down syndrome group; F, females, M, males. *p<0.05, **p<0.01, ***p<0.001.

Subsequent Bayesian analyses indicated strong evidence in favour of reduced calretinin and calbindin proportions (BF10 = 11.41 and 19.43, respectively) and moderate evidence in favour of reduced parvalbumin proportion (BF10 = 5.36), over a null hypothesis of no difference in proportions between controls and individuals with Down syndrome.

### Multiple Regression

Backward regression analysis identified a model containing diagnosis and age as the best model predicting normalised calretinin expression (adjusted *R*^*2*^ = 0.36, *F*(2,24) = 8.20, p_c_ = 0.006). In contrast, models containing only diagnosis were the best at predicting parvalbumin (adjusted *R*^*2*^ = 0.29, *F*(1, 25) = 11.71, *p*_*c*_ = 0.006) and calbindin expression (adjusted *R*^*2*^ = 0.47, *F*(1, 24) = 23.16, *p*_c_ < 0.001). A model containing Aβ42 neared significance as the best predictor of increasing neuronal density (adjusted *R*^*2*^ = 0.12, *F*(1,25) = 4.45, *p*_*c*_ = 0.056).

Bayesian regression analyses were carried out using diagnosis, age, and Aβ42 as predictors of normalized calretinin, parvalbumin, and calbindin expression. For calretinin, there was strong evidence that the model with diagnosis and age explained a large amount of the variance, *R*^*2*^ = 0.41, (BF_10_ = 21.47). Diagnosis provided moderate evidence (β_1_ =-0.17, BF_inclusion_ = 6.25) while Age (β_2_ = - 0.003, BF_inclusion_ = 1.84) and Aβ42 (β_3_ = -0.012, BF_inclusion_ = 1.84) provided only anecdotal evidence for inclusion in the model. For parvalbumin, there was strong evidence that the model with diagnosis alone explained a large amount of variance *R*^*2*^ = 0.32 (BF_10_ = 16.70). Diagnosis provided moderate evidence for inclusion in the model (β_1_ =-0.21, BF_inclusion_ = 7.11). In contrast, both age (β_2_ =-0.00036, BF_inclusion_ = 0.57) and Aβ42 (β_3_ =0.003, BF_inclusion_ = 0.59) provided anecdotal evidence for exclusion from the model. Finally, for calbindin, there was strong evidence that the model with diagnosis alone explained a large amount of variance *R*^*2*^ = 0.49 (BF_10_ = 290). Diagnosis provided strong evidence for inclusion in the model (β_1_ =-0.29, BF_inclusion_ = 25.85). In contrast, both age (β_2_ =-0.00094, BF_inclusion_ = 0.57) and Aβ42 (β_3_ =-0.014, BF_inclusion_ = 0.57) provided anecdotal evidence for exclusion from the model.

## Discussion

Despite the longstanding association between the medial diencephalon and memory, there has been surprisingly little research into the status of this brain region in individuals with Down syndrome. In the current study we assessed the anteroventral thalamic nucleus (AV) in post-mortem tissue from 15 individuals with Down syndrome (DS group). In addition to measuring surface area and neuronal density, we also measured calcium-binding protein immunoreactivity; calcium-binding proteins play an important role in calcium homeostasis and intracellular signalling and have been linked to neurodegeneration (Kelemen and Szilágyi, 2021).

As expected, we found higher levels of amyloid markers in the DS group, most notably in total Aβ burden (as measured by Aβ-4G8) and diffuse Aβ plaques (as measured by Aβ42 expression). Expression of Aβ40-positive mature plaques was generally low with near zero expression in 13 of 15 samples from individuals with Down syndrome. These findings are in line with previous reports showing that Aβ42 plaques are more prevalent than Aβ40 plaques across all ages in Down syndrome (Head et al., 2016). Previous studies have found less pronounced tau expression in the anterior thalamic nuclei in Alzheimer’s disease than other regions (Rüb et al., 2016, Braak and Braak, 1991, Viney et al., 2022). Consistent with this, there was very little tau pathology in most of the cases we examined across groups with near zero levels of tau in 12 of the 15 cases from the DS group. While there were too few cases with high tau expression to draw any clear conclusions, there did not seem to be an association with age as the individual with the highest tau levels was 49 years old, but that case also had the lowest neuron density, suggesting neurons may be particularly vulnerable to high levels of tau in the AV. The heterogeneity in both tau and amyloid expression in younger cases suggests some individuals may have better protection against earlier Alzheimer’s pathology (Maxwell et al., 2021).

Astroglial, but not oligodendrocyte, density was significantly increased in the AV in the DS group. In addition, there was an increase in neuron density in the AV of individuals with Down syndrome, although there was far greater variability in the DS group with some showing very low neuron density and others showing increased neuron density. The overall pattern of increased neuronal and glial density may appear to contradict our previous findings (Perry et al., 2019), where we found reduced total neuron and glia counts. However, the increased neuron density in the current study likely reflects increased packing due to a reduction in AV volume. Due to sample availability, we were not able to acquire total volume measures, as we were in our previous study (Perry et al., 2019), but we acquired a surface area measure for AV for each case to provide a relative approximation of size. The surface area of AV was approximately 37% smaller in the DS group than controls, therefore, even with preserved or increased neuronal density, there are likely fewer neurons overall in the AV in the DS group. Only a few studies have quantitatively assessed the anterior thalamic nuclei in Alzheimer’s disease (De Simone et al., 2024) and even fewer have carried out neuronal counts but, from the studies that are available, there does not seem to be a significant reduction in either volume or neuronal counts in this region using post-mortem tissue (Hornberger et al., 2012, Xuereb et al., 1991), suggesting that the AV may be more impacted in Down syndrome than Alzheimer’s disease alone.

A further aim of the current study was to assess calcium-binding proteins in the AV in Down syndrome. The proportion of AV neurons immunopositive for calbindin, calretinin and parvalbumin was significantly reduced in the DS group, by 70%, 49% and 61%, respectively. Furthermore, we found strong evidence in favour of reduced proportions of calretinin and calbindin positive cells and moderate evidence in favour of reduced proportions of parvalbumin positive cells, according to the Bayesian framework. A previous study had reported a reduction in both calbindin and parvalbumin in the cortices of four older individuals with Down syndrome (Kobayashi et al., 1990). For calbindin, they found a 38% and 28% reduction in immunoreactive neurons in frontal and temporal cortices, respectively. The reported reduction in parvalbumin was slightly less than calbindin, with a 27% decrease in frontal cortex and 22% decrease in temporal cortex. Consistent with these cortical findings (Kobayashi et al., 1990), we also found a greater decrease in calbindin than parvalbumin. However, the relative reduction for both parvalbumin and calbindin appears greater in AV than in the cortex, especially considering our sample population also included younger individuals. In our data set, there was increased neuronal density in the DS group but reduced calcium-binding protein immunoreactivity; this would suggest a downregulation in expression rather than a loss of neurons. However, we cannot tell from this data set whether there are inherent differences in these neuron populations in Down syndrome, i.e., the expression profile is different from birth, or whether downregulation occurs due to pathophysiological conditions or, for example, changes in BDNF-levels, which are altered in Down syndrome and affect calcium-binding protein expression (Fairless et al., 2019).

Parvalbumin immunoreactivity has been previously analysed in human AV. One study found 53-58% of total neurons to be immunoreactive for parvalbumin (Dixon et al., 2000) while another found the percentage of parvalbumin positive cells to range from 55-63% (Danos et al., 1998). In our study, we found approximately 45% of cells to be immunoreactive for parvalbumin in the control group. The methodology and samples are very different across studies, but it seems a consistent finding that approximately half the neurons in AV are immunoreactive for parvalbumin.

Reductions in parvalbumin, calretinin, and calbindin immunoreactivity have been reported in Alzheimer’s disease (Zallo et al., 2018, Brady and Mufson, 1997, Kook et al., 2014b), however, these studies have typically focused on the hippocampus and frontal cortex; to our knowledge, no study has specifically assessed these markers in the anterior thalamic nuclei in Alzheimer’s disease. It is, therefore, not clear whether the reduction in calcium-binding proteins in the DS group simply reflects a sensitivity to Alzheimer-related pathology or whether there are underlying changes to the cell populations in Down syndrome that co-occur with, or even pre-date, amyloid changes. There did not seem to be a clear relationship between amyloid or tau and calcium-binding protein expression but there were cases with very low amyloid and tau expression that also had very low levels of calbindin, parvalbumin and calretinin expression, consistent with the calcium-binding protein changes being found irrespective of amyloid status. In support of this, according to our most parsimonious frequentist and Bayesian regression models, Aβ42, the most pathological form of these Aβ species (Bodani et al., 2015, Iwatsubo et al., 1995), did not predict expression of any calcium-binding protein. Down syndrome alone was the best predictor of calcium-binding protein expression. In comparison, advancing age and Aβ42 deposition provided only some evidence in favour of predicting calretinin expression, but some evidence to be excluded as predictors of parvalbumin and calbindin expression. It is possible that biological age could be a stronger predictor than the chronological age used in this study. Recently, others have proposed mathematical models to predict a brain’s biological age based on MRI derived measures of cortical thickness, cortical surface area, and subcortical volumes (Liem et al., 2017, Salih et al., 2023). Future human tissue studies could estimate brain age using stereological assessment of cortical and subcortical volumes. While there does not seem to be a clear association between Alzheimer markers and calcium-binding protein expression in this current data set, it is possible that earlier changes, for example Aβ and/or tau seeding (Ye et al., 2017, Condello et al., 2022), may show clearer links. A reduction in calcium-binding proteins would disrupt calcium homeostasis, and a loss of calbindin in a rodent model of Alzheimer’s disease was linked to increased pathogenesis (Kook et al., 2014a). It is, therefore, possible that an early loss of calbindin could exacerbate Alzheimer-related pathology in Down syndrome, given calbindin shows the greatest reduction in the present study.

While there are few studies on the thalamus in Down syndrome, a recent study assessed three thalamic nuclei in tissue from foetuses with Down syndrome: mediodorsal nucleus, centromedian nucleus and parafasciular nucleus (Stagni et al., 2020). A similar pattern was found across all nuclei, with a reduction in neuron and glia density but no difference in surface area, consistent with an overall reduction in total neurons in these thalamic regions. There was high expression of calretinin in the mediodorsal thalamic nucleus, with approximately 80% of cells staining positive for calretinin but there was no difference in expression between the control and Down syndrome group (Stagni et al., 2020). This might suggest that expression levels are generally comparable across individuals with Down syndrome and controls in early life, with downregulation only occurring with increasing age. Alternatively, there may be variability across thalamic nuclei and the age at which changes occur, which might be expected given the different expression profiles and functions across thalamic nuclei (Stagni et al., 2020, Morel et al., 1997, Münkle et al., 2000, Wolff and Vann, 2019). Similarly, a stereological study of the superior temporal gyrus of four males with Down syndrome aged 19-33 years found reduced neurons but preserved parvalbumin and a non-significant reduction in calretinin-positive neurons (Giffin-Rao et al., 2022). This reinforces the importance of future studies into the anterior thalamic nuclei in earlier age groups to identify whether there is downregulation in calcium-binding protein in early life or if changes only occur after adolescence. If there is a downregulation of calcium-binding protein expression after adolescence, then finding ways to preserve the expression of these proteins may help protect against subsequent pathogenesis (Hijazi et al., 2020).

Unlike other cortical and allocoritcal regions, the majority of calcium-binding positive neurons in the primate anterior thalamus are projection neurons rather than interneurons (Jones and Hendry, 1989, Dixon et al., 2000). Although calcium-binding protein expressing neurons are thought to largely comprise distinct neuronal populations, it is possible that these proteins are co-expressed in some anterior thalamic neurons. In support of this, neuronal co-expression of these proteins has been reported in the human visual and temporal cortex regions (del Río and DeFelipe, 1997, Leuba and Saini, 1997) and in the anterior thalamus of guinea pigs (Zakowski et al., 2013). We were not able to assess colocalization in the current study, however, the total proportion of calcium-binding protein positive neurons was greater than one among control samples but less than one among samples from people with Down syndrome. This suggests some colocalization of neuronal calcium-binding protein expression may be occurring, especially among control samples, highlighting the need for future studies to examine the co-expression of calcium-binding proteins in the human anterior thalamus.

The current study used digital pathology methods. This enabled objective assessment of all the stained objects in the AV on each section. While this method to identify and quantify cells and for measurements of surface area is suitable for identifying relative numbers between groups, it cannot estimate the true number of cells or surface area in these samples. This can only be achieved with stereological assessment which requires access to all the tissue from the region of interest. Much of the tissue from current study is historical, with portions of the anterior thalamus having been used in other studies. However, despite the lack of stereological assessments, the data still provides valuable insights to changes in the anterior thalamus of younger and older adults with Down syndrome given the rarity of this tissue and the lack of knowledge of the anterior thalamus in Down syndrome.

Although, there was variation in the anteroposterior level assessed across cases, the sections for both the DS and control groups came from a similar range of anteroposterior levels. Furthermore, cell density and calretinin immunoreactivity in the human anterior thalamus is uniform across the anteroposterior extent of the nuclei (Braak and Weinel, 1985, Morel et al., 1997, Fortin et al., 1998), giving additional weight to the findings. In immunohistochemistry studies, the reliability of findings can be influenced by variations in tissue handling and staining and this is especially important when few sections per case are stained. This source of variation was reduced in the current study as samples were sourced from UKBBN banks that collected tissue according to an agreed common standard and all sections for each immunohistochemistry procedure were processed in a single session.

The present study adds to the growing literature linking neuropathology in the medial diencephalon, including anterior thalamic nuclei and mammillary bodies, to various neurological disorders (Milczarek et al., 2023a, Lequin et al., 2021, Meys et al., 2022, Clarke et al., 1994, Connaughton et al., 2023, Danos et al., 1998). Pathology in these brain areas would contribute to memory impairments often accompanying these conditions by affecting both the encoding and consolidation of memories (Milczarek et al., 2023b, Aggleton et al., 2023). In the present study, we found changes in the AV in Down syndrome at a younger age than previously reported and showed for the first time that there is a marked loss of calcium-binding protein immunoreactivity. Noticeably, this reduction in parvalbumin, calretinin and calbindin immunoreactivity was not simply driven by overall neuronal loss but likely reflects a downregulation in expression. These reductions in calcium-binding proteins may increase the vulnerability of cells within this region and potentially exacerbate expression of Alzheimer-related pathology.

## Supporting information

Addition file 1

## Acknowledgements

This research was funded by a grant from the Jerome Lejeune Foundation (# 1899) and a Wellcome Trust Senior Research Fellowship (WT212273/Z/18) awarded to SDV.

Tissue samples were sourced from brain banks in the UK Brain Banks Network. The Manchester Brain Bank, which is part of the Brains for Dementia Research Initiative, jointly funded by Alzheimer’s Society and Alzheimer’s Research UK. The Cambridge Brain Bank is supported by the NIHR Cambridge Biomedical Research Centre (NIHR203312). We acknowledge the Oxford Brain Bank, supported by Brains Dementia Research (BDR) (Alzheimer Society and Alzheimer Research UK) and the National Institute for Health Research (NIHR) Oxford Biomedical Research Centre (BRC). Tissue samples were provided by The London Neurodegenerative Diseases Brain Bank at King’s College London. The brain bank receives funding as part of the Brains for Dementia Research programme, jointly funded by Alzheimer’s Research UK and the Alzheimer’s Society.

## List of abbreviations

AV: anteroventral thalamic nucleus
Aβ: beta-amyloid
BF_10_: Bayes Factor
DAB: 3,3’-diaminobenzidine
DS: Down syndrome.

