## Supplementary material for "Reduction in neurons immunoreactive for parvalbumin, calretinin and calbindin in the anteroventral thalamic nuclei of individuals with Down syndrome": Addition file 1

**Additional file 1**. Supplementary tables 1 through 2.

**Supplementary table 1**. Demographic and biological data for each case.

| UKBBN | Brain bank | Diagnosis | Age | Sex | PMD(h) | Brain weight (g) |
| --- | --- | --- | --- | --- | --- | --- |
| BBN110.30102 | CAM | Control | 38 | M | 84 | 1517 |
| BBN_15790 | LON | Control | 40 | M | 40 | NA |
| BBN002.30139 | LON | Control | 47 | F | 26 | 1087 |
| BBN_17100 | LON | Control | 48 | M | 60 | NA |
| BBN110.28668 | CAM | Control | 60 | F | 60 | 1090 |
| BBN002.29402 | LON | Control | 60 | F | 33 | 1168 |
| BBN110.28899 | CAM | Control | 61 | F | 84 | 1072 |
| BBN110.35427 | CAM | Control | 63 | M | 83 | 1513 |
| BBN_16251 | LON | Control | 66 | M | 52 | NA |
| BBN002.28414 | LON | Control | 68 | F | 58.5 | NA |
| BBN_24559 | LON | Control | 68 | F | 47 | NA |
| BBN_16993 | LON | Control | 70 | M | 23 | NA |
| BBN004.26633 | OXF | DS | 22 | M | 96 | NA |
| BBN004.26750 | OXF | DS | 35 | M | NA | NA |
| BBN004.26635 | OXF | DS | 36 | M | 72 | NA |
| BBN002.28703 | LON | DS | 37 | M | 48 | 1238 |
| BBN004.26644 | OXF | DS | 39 | F | 95 | NA |
| BBN_17198 | LON | DS | 42 | F | 100 | NA |
| BBN004.26630 | OXF | DS | 42 | F | NA | NA |
| BBN_17186 | LON | DS | 47 | F | NA | NA |
| BBN_17189 | LON | DS | 47 | M | 24 | NA |
| BBN_16835 | LON | DS | 60 | M | 60 | NA |
| BBN_2990 | MAN | DS | 60 | F | NA | 950 |
| BBN_2984 | MAN | DS | 61 | M | NA | 1018 |
| BBN_3439 | MAN | DS | 62 | M | NA | NA |
| BBN_2967 | MAN | DS | 62 | F | NA | 960 |
| BBN_2975 | MAN | DS | 65 | M | NA | NA |

*Note.* Cam, Cambridge Brain Bank; Lon, London Neurodegenerative Disease Brain Bank; Man, Manchester Brain Bank; Oxf, Oxford Brain Bank; PMD, Post-mortem delay.

**Supplementary table 2**. Performance (%) of the cell classification models.

|  |  | Neurons | Astroglia | Oligo | Microglia | Calretinin | Calbindin | Parvalbumin |
| --- | --- | --- | --- | --- | --- | --- | --- | --- |
| Accuracy | C  DS | 98.8  99 | 95.1  97.8 | 95.3  97.8 | 98.3  99.7 | 89.8  89.7 | 90.2  92.2 | 88.9  90.8 |
| Precision | C  DS | 97.1  92.1 | 93.5  96.7 | 94.5  96.7 | 86.4  90.9 | 89.9  89.9 | 91.9  94.5 | 94.3  98.1 |
| Recall | C  DS | 89.2  97.2 | 97.1  98.2 | 91.8  98.8 | 82.6  90.9 | 95.2  94 | 96.5  95 | 91.2  91.3 |

*Note*: Accuracy is the number of cells correctly classified out of all cells present. Precision is the number of cells belonging to the correct classification out of all the samples that were predicted to be of that classification. Recall is the number of cells correctly classified out of all samples that belong to that classification. The formula for each calculation is provided below. TP, true positive; TN, true negative; FP, false positive; FN, false negative. C, control; DS, Down syndrome.

$$Accuracy=\frac{TP+TN}{TP+FP+TN+FN}$$

$$Precision=\frac{TP}{TP+FP}$$

$$Recall=\frac{TP}{TP+FN}$$

**Supplementary table 3.** Peason’s correlations with post-mortem delay

| **Measure** | **Pearson’s *r*** | ***p*** |
| --- | --- | --- |
| AV area | 0.052 | 0.83 |
| Neuron density | 0.070 | 0.78 |
| Total glia density | -0.179 | 0.46 |
| Astroglia density | -0.008 | 0.97 |
| Oligo density | -0.279 | 0.25 |
| Microglia density | 0.135 | 0.58 |
| Aβ-4G8 | -0.288 | 0.23 |
| Aβ42 | -0.038 | 0.88 |
| Aβ40 | -0.352 | 0.14 |
| Tau AT8 | -0.050 | 0.85 |
| Calretinin proportion | -0.268 | 0.27 |
| Parvalbumin proportion | -0.091 | 0.71 |
| Calbindin proportion | -0.100 | 0.69 |
| Calretinin density | -0.447 | 0.06 |
| Parvalbumin density | -0.135 | 0.58 |
| Calbindin density | -0.190 | 0.45 |

*Note:* *p*-values are uncorrected for multiple comparisons.
