## Additional file 2 for "Reduction in neurons immunoreactive for parvalbumin, calretinin and calbindin in the anteroventral thalamic nuclei of individuals with Down syndrome"

**Additional file 2**: Supplementary tables 3 through 14.

| **Supplementary table 3**. Calretinin proportion regression model summary. | | | | | | | | | | | | | | |
| --- | --- | --- | --- | --- | --- | --- | --- | --- | --- | --- | --- | --- | --- | --- |
|  | | | | | | | | | | **Durbin-Watson** | | | | |
| **Model** | | **R** | | **R²** | | **Adjusted R²** | | **RMSE** | | **Autocorrelation** | | **Statistic** | | **p** |
| 1 |  | 0.637 |  | 0.406 |  | 0.329 |  | 0.161 |  | -0.138 |  | 2.119 |  | 0.975 |
| 2 |  | 0.637 |  | 0.406 |  | 0.356 |  | 0.158 |  | -0.125 |  | 2.095 |  | 0.955 |

*Note*: Model 2, the most parsimonious model had acceptable independence of errors, Durbin-Watson = 2.10. RMSE, root mean squared error.

| **Supplementary table 4.** Calretinin proportion regression model ANOVA. | | | | | | | | | | | | |
| --- | --- | --- | --- | --- | --- | --- | --- | --- | --- | --- | --- | --- |
| **Model** | |  | | **Sum of Squares** | | **df** | | **Mean Square** | | **F** | | **p** |
| 1 |  | Regression |  | 0.409 |  | 3 |  | 0.136 |  | 5.246 |  | 0.007 |
|  |  | Residual |  | 0.598 |  | 23 |  | 0.026 |  |  |  |  |
|  |  | Total |  | 1.008 |  | 26 |  |  |  |  |  |  |
| 2 |  | Regression |  | 0.409 |  | 2 |  | 0.205 |  | 8.202 |  | 0.002 |
|  |  | Residual |  | 0.599 |  | 24 |  | 0.025 |  |  |  |  |
|  |  | Total |  | 1.008 |  | 26 |  |  |  |  |  |  |

*Note*: The *p* values are not corrected for multiple comparisons.

| **Supplementary table 5.** Calretinin proportion regression model coefficients. | | | | | | | | | | | | | | | | |
| --- | --- | --- | --- | --- | --- | --- | --- | --- | --- | --- | --- | --- | --- | --- | --- | --- |
|  | | | | | | | | | | | | | | **Collinearity Statistics** | | |
| **Model** | |  | | **Unstandardized (*b*)** | | **Standard Error** | | **Standardized (β)** | | **t** | | **p** | | **Tolerance** | | **VIF** |
| 1 |  | (Intercept) |  | 0.736 |  | 0.183 |  |  |  | 4.032 |  | 5.197×10^-4^ |  |  |  |  |
|  |  | Age |  | -0.005 |  | 0.003 |  | -0.349 |  | -1.703 |  | 0.102 |  | 0.614 |  | 1.628 |
|  |  | Aβ42 |  | -0.005 |  | 0.046 |  | -0.026 |  | -0.108 |  | 0.915 |  | 0.448 |  | 2.230 |
|  |  | Diagnosis |  | -0.255 |  | 0.099 |  | -0.657 |  | -2.574 |  | 0.017 |  | 0.396 |  | 2.523 |
| 2 |  | (Intercept) |  | 0.746 |  | 0.154 |  |  |  | 4.842 |  | 6.202×10^-5^ |  |  |  |  |
|  |  | Age |  | -0.005 |  | 0.003 |  | -0.361 |  | -2.128 |  | 0.044 |  | 0.860 |  | 1.163 |
|  |  | Diagnosis |  | -0.263 |  | 0.066 |  | -0.677 |  | -3.992 |  | 5.380×10^-4^ |  | 0.860 |  | 1.163 |

*Note*: The variance inflation factor values for each predictor variable were less than 2.53 indicating there were not concerns with multicollinearity. The regression equation for the most parsimonious model is **calretinin proportion = 0.746 + (-0.005 x age in years) + (-0.263 x diagnosis)** where a diagnosis of Down syndrome = 1 and control = 0.

| **Supplementary table 6**. Parvalbumin proportion regression model summary. | | | | | | | | | | | | | | |
| --- | --- | --- | --- | --- | --- | --- | --- | --- | --- | --- | --- | --- | --- | --- |
|  | | | | | | | | | | **Durbin-Watson** | | | | |
| **Model** | | **R** | | **R²** | | **Adjusted R²** | | **RMSE** | | **Autocorrelation** | | **Statistic** | | **p** |
| 1 |  | 0.573 |  | 0.328 |  | 0.240 |  | 0.216 |  | 0.035 |  | 1.927 |  | 0.584 |
| 2 |  | 0.567 |  | 0.322 |  | 0.265 |  | 0.213 |  | -0.028 |  | 2.052 |  | 0.865 |
| 3 |  | 0.565 |  | 0.319 |  | 0.292 |  | 0.209 |  | -0.002 |  | 1.999 |  | 0.841 |

*Note*: Model 3, the most parsimonious model had acceptable independence of errors, Durbin-Watson = 2.00. RMSE, root mean squared error.

| **Supplementary table 7.** Parvalbumin proportion regression model ANOVA. | | | | | | | | | | | | |
| --- | --- | --- | --- | --- | --- | --- | --- | --- | --- | --- | --- | --- |
| **Model** | |  | | **Sum of Squares** | | **df** | | **Mean Square** | | **F** | | **p** |
| 1 |  | Regression |  | 0.526 |  | 3 |  | 0.175 |  | 3.743 |  | 0.025 |
|  |  | Residual |  | 1.078 |  | 23 |  | 0.047 |  |  |  |  |
|  |  | Total |  | 1.604 |  | 26 |  |  |  |  |  |  |
| 2 |  | Regression |  | 0.516 |  | 2 |  | 0.258 |  | 5.695 |  | 0.009 |
|  |  | Residual |  | 1.088 |  | 24 |  | 0.045 |  |  |  |  |
|  |  | Total |  | 1.604 |  | 26 |  |  |  |  |  |  |
| 3 |  | Regression |  | 0.512 |  | 1 |  | 0.512 |  | 11.707 |  | 0.002 |
|  |  | Residual |  | 1.092 |  | 25 |  | 0.044 |  |  |  |  |
|  |  | Total |  | 1.604 |  | 26 |  |  |  |  |  |  |

*Note*: The *p* values are not corrected for multiple comparisons.

| **Supplementary table 8.** Parvalbumin proportion regression model coefficients. | | | | | | | | | | | | | | | | |
| --- | --- | --- | --- | --- | --- | --- | --- | --- | --- | --- | --- | --- | --- | --- | --- | --- |
|  | | | | | | | | | | | | | | **Collinearity Statistics** | | |
| **Model** | |  | | **Unstandardized (*b*)** | | **Standard Error** | | **Standardized (β)** | | **t** | | **p** | | **Tolerance** | | **VIF** |
| 1 |  | (Intercept) |  | 0.572 |  | 0.245 |  |  |  | 2.333 |  | 0.029 |  |  |  |  |
|  |  | Diagnosis |  | -0.333 |  | 0.133 |  | -0.679 |  | -2.500 |  | 0.020 |  | 0.396 |  | 2.523 |
|  |  | Age |  | -0.002 |  | 0.004 |  | -0.112 |  | -0.513 |  | 0.613 |  | 0.614 |  | 1.628 |
|  |  | Aβ42 |  | 0.028 |  | 0.062 |  | 0.118 |  | 0.462 |  | 0.648 |  | 0.448 |  | 2.230 |
| 2 |  | (Intercept) |  | 0.514 |  | 0.208 |  |  |  | 2.476 |  | 0.021 |  |  |  |  |
|  |  | Diagnosis |  | -0.288 |  | 0.089 |  | -0.586 |  | -3.236 |  | 0.004 |  | 0.860 |  | 1.163 |
|  |  | Age |  | -0.001 |  | 0.003 |  | -0.058 |  | -0.321 |  | 0.751 |  | 0.860 |  | 1.163 |
| 3 |  | (Intercept) |  | 0.451 |  | 0.060 |  |  |  | 7.472 |  | 7.972×10^-8^ |  |  |  |  |
|  |  | Diagnosis |  | -0.277 |  | 0.081 |  | -0.565 |  | -3.422 |  | 0.002 |  | 1.000 |  | 1.000 |

*Note*: The variance inflation factor values for each predictor variable were less than 2.53 indicating there were not concerns with multicollinearity. The regression equation for the most parsimonious model is **parvalbumin proportion = 0.451 + (-0.277 x diagnosis)** where a diagnosis of Down syndrome = 1 and control = 0.

| **Supplementary table 9**. Calbindin proportion regression model summary. | | | | | | | | | | | | | | | | | | | |
| --- | --- | --- | --- | --- | --- | --- | --- | --- | --- | --- | --- | --- | --- | --- | --- | --- | --- | --- | --- |
|  | | | | | | | | | | | | **Durbin-Watson** | | | | | | | |
| **Model** | | **R** | | **R²** | **Adjusted R²** | | | **RMSE** | | | **Autocorrelation** | | | | **Statistic** | | | **p** | |
| 1 |  | 0.716 |  | 0.512 |  | 0.446 |  | | 0.197 |  | | | -0.162 |  | | 2.156 |  | | 0.964 |
| 2 |  | 0.705 |  | 0.497 |  | 0.453 |  | | 0.196 |  | | | -0.070 |  | | 1.971 |  | | 0.690 |
| 3 |  | 0.701 |  | 0.491 |  | 0.470 |  | | 0.193 |  | | | -0.043 |  | | 1.912 |  | | 0.665 |

*Note*: Model 3, the most parsimonious model had acceptable independence of errors, Durbin-Watson = 1.91. RMSE, root mean squared error.

| **Supplementary table 10.** Calbindin proportion regression model ANOVA. | | | | | | | | | | | | |
| --- | --- | --- | --- | --- | --- | --- | --- | --- | --- | --- | --- | --- |
| **Model** | |  | | **Sum of Squares** | | **df** | | **Mean Square** | | **F** | | **p** |
| 1 |  | Regression |  | 0.896 |  | 3 |  | 0.299 |  | 7.705 |  | 0.001 |
|  |  | Residual |  | 0.852 |  | 22 |  | 0.039 |  |  |  |  |
|  |  | Total |  | 1.748 |  | 25 |  |  |  |  |  |  |
| 2 |  | Regression |  | 0.868 |  | 2 |  | 0.434 |  | 11.348 |  | 3.725×10^-4^ |
|  |  | Residual |  | 0.880 |  | 23 |  | 0.038 |  |  |  |  |
|  |  | Total |  | 1.748 |  | 25 |  |  |  |  |  |  |
| 3 |  | Regression |  | 0.858 |  | 1 |  | 0.858 |  | 23.160 |  | 6.690×10^-5^ |
|  |  | Residual |  | 0.890 |  | 24 |  | 0.037 |  |  |  |  |
|  |  | Total |  | 1.748 |  | 25 |  |  |  |  |  |  |

*Note*: The *p* values are not corrected for multiple comparisons.

**Supplementary table 11.** Calbindin proportion regression model coefficients.

|  | | | | | | | | | | | | | | | | | | | | |
| --- | --- | --- | --- | --- | --- | --- | --- | --- | --- | --- | --- | --- | --- | --- | --- | --- | --- | --- | --- | --- |
|  | | | | | | | | | | | | | | | | | | **Collinearity Statistics** | | |
| **Model** | |  | | **Unstandardized (*b*)** | | | **Standard Error** | | | **Standardized (β)** | | | **t** | | | **p** | | **Tolerance** | | **VIF** |
| 1 |  | (Intercept) |  | | 0.334 |  | | 0.226 |  | |  |  | | 1.476 |  | 0.154 |  |  |  |  |
|  |  | Diagnosis |  | | -0.280 |  | | 0.121 |  | | -0.533 |  | | -2.309 |  | 0.031 |  | 0.416 |  | 2.402 |
|  |  | Age |  | | 0.003 |  | | 0.004 |  | | 0.166 |  | | 0.881 |  | 0.388 |  | 0.622 |  | 1.609 |
|  |  | Aβ42 |  | | -0.048 |  | | 0.057 |  | | -0.187 |  | | -0.841 |  | 0.410 |  | 0.448 |  | 2.234 |
| 2 |  | (Intercept) |  | | 0.433 |  | | 0.192 |  | |  |  | | 2.252 |  | 0.034 |  |  |  |  |
|  |  | Diagnosis |  | | -0.354 |  | | 0.082 |  | | -0.674 |  | | -4.296 |  | 2.694×10^-4^ |  | 0.888 |  | 1.126 |
|  |  | Age |  | | 0.002 |  | | 0.003 |  | | 0.079 |  | | 0.505 |  | 0.618 |  | 0.888 |  | 1.126 |
| 3 |  | (Intercept) |  | | 0.525 |  | | 0.058 |  | |  |  | | 9.048 |  | 3.338×10^-9^ |  |  |  |  |
|  |  | Diagnosis |  | | -0.368 |  | | 0.076 |  | | -0.701 |  | | -4.812 |  | 6.690×10^-5^ |  | 1.000 |  | 1.000 |

*Note:* The variance inflation factor values for each predictor variable were less than 2.41 indicating there were not concerns with multicollinearity. The regression equation for the most parsimonious model is **calbindin proportion = 0.525 + (-0.368 x diagnosis)** where a diagnosis of Down syndrome = 1 and control = 0.

| **Supplementary table 12**. Neuron density regression model summary. | | | | | | | | | | | | | | |
| --- | --- | --- | --- | --- | --- | --- | --- | --- | --- | --- | --- | --- | --- | --- |
|  | | | | | | | | | | **Durbin-Watson** | | | | |
| **Model** | | **R** | | **R²** | | **Adjusted R²** | | **RMSE** | | **Autocorrelation** | | **Statistic** | | **p** |
| 1 |  | 0.430 |  | 0.185 |  | 0.078 |  | 59.171 |  | 0.008 |  | 1.971 |  | 0.667 |
| 2 |  | 0.429 |  | 0.184 |  | 0.116 |  | 57.953 |  | 0.003 |  | 1.981 |  | 0.748 |
| 3 |  | 0.389 |  | 0.151 |  | 0.117 |  | 57.904 |  | -0.083 |  | 2.148 |  | 0.774 |

*Note*: Model 3, the most parsimonious model had acceptable independence of errors, Durbin-Watson = 2.15. RMSE, root mean squared error.

| **Supplementary table 13.** Neuron density regression model ANOVA. | | | | | | | | | | | | | | |
| --- | --- | --- | --- | --- | --- | --- | --- | --- | --- | --- | --- | --- | --- | --- |
| **Model** | |  | **Sum of Squares** | | | **df** | | **Mean Square** | | | **F** | | **p** | |
| 1 |  | Regression |  | 18223.883 |  | | 3 |  | 6074.628 |  | | 1.735 |  | 0.188 |
|  |  | Residual |  | 80527.877 |  | | 23 |  | 3501.212 |  | |  |  |  |
|  |  | Total |  | 98751.759 |  | | 26 |  |  |  | |  |  |  |
| 2 |  | Regression |  | 18145.592 |  | | 2 |  | 9072.796 |  | | 2.701 |  | 0.087 |
|  |  | Residual |  | 80606.167 |  | | 24 |  | 3358.590 |  | |  |  |  |
|  |  | Total |  | 98751.759 |  | | 26 |  |  |  | |  |  |  |
| 3 |  | Regression |  | 14930.456 |  | | 1 |  | 14930.456 |  | | 4.453 |  | 0.045 |
|  |  | Residual |  | 83821.303 |  | | 25 |  | 3352.852 |  | |  |  |  |
|  |  | Total |  | 98751.759 |  | | 26 |  |  |  | |  |  |  |

*Note*: The *p* values are not corrected for multiple comparisons.

| **Supplementary table 14.** Neuron density regression model coefficients. | | | | | | | | | | | | | | | | |
| --- | --- | --- | --- | --- | --- | --- | --- | --- | --- | --- | --- | --- | --- | --- | --- | --- |
|  | | | | | | | | | | | | | | **Collinearity Statistics** | | |
| **Model** | |  | | **Unstandardized (b)** | | **Standard Error** | | **Standardized (β)** | | **t** | | **p** | | **Tolerance** | | **VIF** |
| 1 |  | (Intercept) |  | 91.560 |  | 67.004 |  |  |  | 1.366 |  | 0.185 |  |  |  |  |
|  |  | Diagnosis |  | 31.022 |  | 36.398 |  | 0.255 |  | 0.852 |  | 0.403 |  | 0.396 |  | 2.523 |
|  |  | Age |  | 0.170 |  | 1.137 |  | 0.036 |  | 0.150 |  | 0.882 |  | 0.614 |  | 1.628 |
|  |  | Aβ42 |  | 13.603 |  | 16.822 |  | 0.227 |  | 0.809 |  | 0.427 |  | 0.448 |  | 2.230 |
| 2 |  | (Intercept) |  | 101.248 |  | 16.746 |  |  |  | 6.046 |  | 3.044×10^-6^ |  |  |  |  |
|  |  | Diagnosis |  | 27.719 |  | 28.330 |  | 0.228 |  | 0.978 |  | 0.338 |  | 0.628 |  | 1.593 |
|  |  | Aβ42 |  | 14.948 |  | 13.925 |  | 0.250 |  | 1.073 |  | 0.294 |  | 0.628 |  | 1.593 |
| 3 |  | (Intercept) |  | 110.467 |  | 13.832 |  |  |  | 7.987 |  | 2.423×10^-8^ |  |  |  |  |
|  |  | Aβ42 |  | 23.261 |  | 11.023 |  | 0.389 |  | 2.110 |  | 0.045 |  | 1.000 |  | 1.000 |

*Note*: The variance inflation factor values for each predictor variable were less than 2.53 indicating there were not concerns with multicollinearity. The regression equation for the most parsimonious model is **neuron density = 110.47 + (23.26 x percent area occupied by Aβ42)**.
