## Additional file 3 for "Reduction in neurons immunoreactive for parvalbumin, calretinin and calbindin in the anteroventral thalamic nuclei of individuals with Down syndrome"

**Additional file 3**. Supplementary figures 1 through 3.

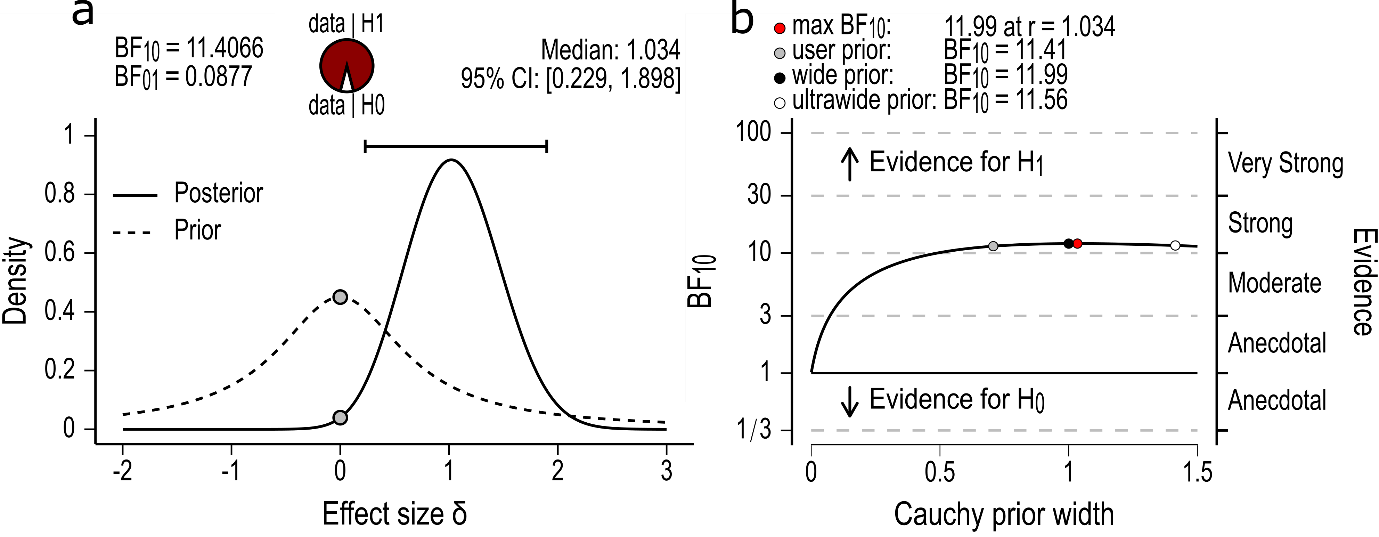

**Supplementary Fig. 1.** Calretinin proportion Bayesian independent means *t* test for the parameter, standardised effect size (δ). Left panel, (a) The probability wheel on top visualizes the relative evidence the data provide for the null hypothesis (H_0_) in white and the alternative hypothesis (H_1_) in red. The dashed line shows the prior distribution and the solid line the posterior distribution. The two gray dots indicate the prior and posterior density at the test value, δ = 0.0. The higher dot on the prior distribution compared to posterior distribution indicates that the Bayes Factor (BF_10_) supports H_1_. The median and the 95% central credible interval of the posterior distribution are shown in the top right corner. Right panel, (b) The Bayes factor robustness plot. The width (uncertainty) of the prior distribution was set by the user to r = 1/ √2. The plot indicates the BF_10_ for the user specified prior, wide prior (r = 1), and ultrawide prior (r = √2). The BF_10_ is relatively consistent across the range of prior widths indicating that the evidence for H_1_ is robust. Figures produced in JASP. BF_01_, Bayes factor in support of H_0_.

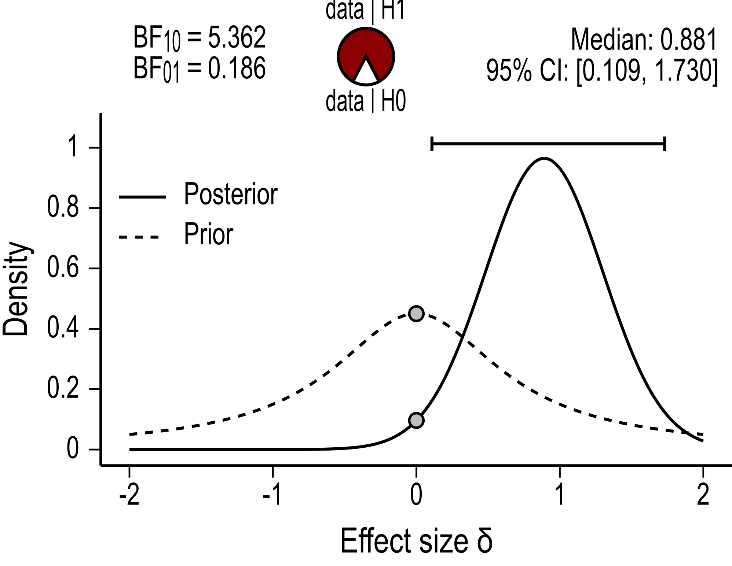

**Supplementary Fig. 2.** Parvalbumin proportion Bayesian Mann-Whitney *U* test for the parameter, standardised effect size (δ). The probability wheel on top visualizes the relative evidence the data provide for the null hypothesis (H_0_) in white and the alternative hypothesis (H_1_) in red. The dashed line shows the prior distribution and the solid line the posterior distribution. The two gray dots indicate the prior and posterior density at the test value, δ = 0.0. The higher dot on the prior distribution compared to posterior distribution indicates that the Bayes Factor (BF_10_) supports H_1_. The median and the 95% central credible interval of the posterior distribution are shown in the top right corner. Note, robustness checks are not available for Mann-Whitney *U* test. The width (uncertainty) of the prior distribution was set by the user to r = 1/ √2. The result is based on Markov Monte Carlo set to five chains of 10, 000 iterations. Figure produced in JASP. BF_01_, Bayes factor in support of H_0_.

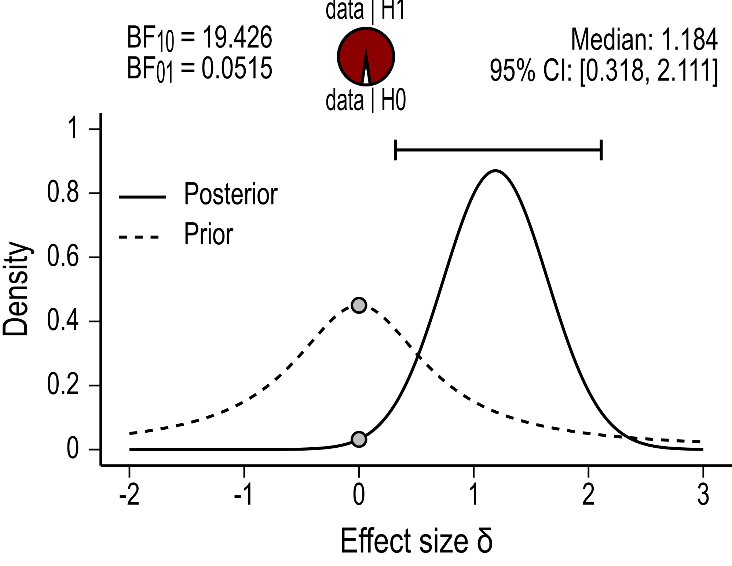

**Supplementary Fig. 3.** Calbindin proportion Bayesian Mann-Whitney *U* test for the parameter, standardised effect size (δ). The probability wheel on top visualizes the relative evidence the data provide for the null hypothesis (H_0_) in white and the alternative hypothesis (H_1_) in red. The dashed line shows the prior distribution and the solid line the posterior distribution. The two gray dots indicate the prior and posterior density at the test value, δ = 0.0. The higher dot on the prior distribution compared to posterior distribution indicates that the Bayes Factor (BF_10_) supports H_1_. The median and the 95% central credible interval of the posterior distribution are shown in the top right corner. The width (uncertainty) of the prior distribution was set by the user to r = 1/ √2. The result is based on Markov Monte Carlo set to five chains of 10, 000 iterations. Note, robustness checks are not available for Mann-Whitney *U* test. Figure produced in JASP. BF_01_, Bayes factor in support of H_0_.
