## Additional file 4 for "Reduction in neurons immunoreactive for parvalbumin, calretinin and calbindin in the anteroventral thalamic nuclei of individuals with Down syndrome"

**Additional file 4**. Supplementary Tables 15 through 23.

| **Supplementary table 15**. Calretinin proportion Bayesian regression model comparison. | | | | | | | | | | |
| --- | --- | --- | --- | --- | --- | --- | --- | --- | --- | --- |
| **Models** | **P(M)** | | **P(M\|data)** | | **BF_M_** | | **BF_10_** | | **R²** | |
| Null model |  | 0.250 |  | 0.040 |  | 0.126 |  | 1.000 |  | 0.000 |
| Diagnosis + Age + Aβ42 |  | 0.250 |  | 0.328 |  | 1.463 |  | 8.108 |  | 0.406 |
| Diagnosis + Age |  | 0.083 |  | 0.289 |  | 4.479 |  | 21.473 |  | 0.406 |
| Diagnosis |  | 0.083 |  | 0.154 |  | 2.002 |  | 11.429 |  | 0.294 |
| Diagnosis + Aβ42 |  | 0.083 |  | 0.091 |  | 1.101 |  | 6.754 |  | 0.331 |
| Aβ42 |  | 0.083 |  | 0.067 |  | 0.784 |  | 4.938 |  | 0.234 |
| Age + Aβ42 |  | 0.083 |  | 0.026 |  | 0.288 |  | 1.896 |  | 0.235 |
| Age |  | 0.083 |  | 0.005 |  | 0.059 |  | 0.399 |  | 0.012 |

*Note:* The two best models explained the same amount of variance (*R*^2^ = 0.41). The most parsimonious model included diagnosis and age as predictors. P(M) = prior model probability; P(M|data) = posterior model probability; BF_M_ = change from prior model odds to posterior model odds; BF_10_ = Bayes factor for each model compared to the null model.

| **Supplementary table 16**. Calretinin proportion posterior summaries of coefficients. | | | | | | | | | | | | | | | | | | | | | |
| --- | --- | --- | --- | --- | --- | --- | --- | --- | --- | --- | --- | --- | --- | --- | --- | --- | --- | --- | --- | --- | --- |
|  | | | | | | | | | | | | | | | | | | **95% Credible Interval** | | | |
| **Coefficient** | | **P(incl)** | **P(excl)** | | **P(incl\|data)** | | | **P(excl\|data)** | | | **BF_inclusion_** | | | **Mean** | | **SD** | | **Lower** | | **Upper** | |
| Intercept |  | 1.000 |  | 0.000 |  | 1.000 |  | | 0.000 |  | | 1.000 |  | | 0.316 |  | 0.032 |  | 0.248 |  | 0.373 |
| Diagnosis |  | 0.500 |  | 0.500 |  | 0.862 |  | | 0.138 |  | | 6.252 |  | | -0.168 |  | 0.099 |  | -0.318 |  | 0.000 |
| Age |  | 0.500 |  | 0.500 |  | 0.648 |  | | 0.352 |  | | 1.841 |  | | -0.003 |  | 0.003 |  | -0.009 |  | 8.309×10^-4^ |
| Aβ42 |  | 0.500 |  | 0.500 |  | 0.511 |  | | 0.489 |  | | 1.044 |  | | -0.012 |  | 0.035 |  | -0.108 |  | 0.043 |

*Note:* P(incl) = prior inclusion probability; P(excl) = prior exclusion probability; P(incl|data) = posterior inclusion probability; P(excl|data) = posterior exclusion probability; BF_inclusion_ = Bayes factor inclusion. The regression equation for the most parsimonious model is **calretinin proportion = 0.32 + (-0.003 x [age – mean age]) + (-0.168*diagnosis)** where a diagnosis of Down syndrome = 1 and control = 0.

**Supplementary table 17**. Descriptive statistics for calretinin Bayesian regression model.

|  | | | | | | |
| --- | --- | --- | --- | --- | --- | --- |
|  | | **N** | | **Mean** | | **SD** |
| Calretinin proportion |  | 27 |  | 0.316 |  | 0.197 |
| Age |  | 27 |  | 52.074 |  | 13.023 |
| Aβ42 |  | 27 |  | 0.743 |  | 1.030 |

| **Supplementary table 18**. Parvalbumin proportion Bayesian regression model comparison. | | | | | | | | | | | |
| --- | --- | --- | --- | --- | --- | --- | --- | --- | --- | --- | --- |
| **Models** | | **P(M)** | **P(M\|data)** | | | **BF_M_** | | **BF_10_** | | **R²** | |
| Null model |  | 0.250 |  | 0.072 |  | | 0.234 |  | 1.000 |  | 0.000 |
| Diagnosis |  | 0.083 |  | 0.402 |  | | 7.395 |  | 16.694 |  | 0.319 |
| Diagnosis + Age + Aβ42 |  | 0.250 |  | 0.194 |  | | 0.721 |  | 2.683 |  | 0.328 |
| Diagnosis + Age |  | 0.083 |  | 0.142 |  | | 1.819 |  | 5.893 |  | 0.322 |
| Diagnosis + Aβ42 |  | 0.083 |  | 0.139 |  | | 1.776 |  | 5.772 |  | 0.320 |
| Aβ42 |  | 0.083 |  | 0.023 |  | | 0.263 |  | 0.968 |  | 0.099 |
| Age + Aβ42 |  | 0.083 |  | 0.017 |  | | 0.186 |  | 0.692 |  | 0.146 |
| Age |  | 0.083 |  | 0.011 |  | | 0.123 |  | 0.458 |  | 0.026 |

*Note:* P(M) = prior model probability; P(M|data) = posterior model probability; BF_M_ = change from prior model odds to posterior model odds; BF_10_ = Bayes factor for each model compared to the null model.

**Supplementary table 19**. Parvalbumin proportion posterior summaries of coefficients.

|  | | | | | | | | | | | | | | | | | | |
| --- | --- | --- | --- | --- | --- | --- | --- | --- | --- | --- | --- | --- | --- | --- | --- | --- | --- | --- |
|  | | | | | | | | | | | | | | | | **95% Credible Interval** | | |
| **Coefficient** | | **P(incl)** | | **P(excl)** | | **P(incl\|data)** | | **P(excl\|data)** | | **BF_inclusion_** | | **Mean** | | **SD** | | **Lower** | | **Upper** |
| Intercept |  | 1.000 |  | 0.000 |  | 1.000 |  | 0.000 |  | 1.000 |  | 0.297 |  | 0.042 |  | 0.208 |  | 0.374 |
| Diagnosis |  | 0.500 |  | 0.500 |  | 0.877 |  | 0.123 |  | 7.114 |  | -0.208 |  | 0.114 |  | -0.400 |  | 0.000 |
| Age |  | 0.500 |  | 0.500 |  | 0.363 |  | 0.637 |  | 0.571 |  | -3.598×10^-4^ |  | 0.002 |  | -0.006 |  | 0.004 |
| Aβ42 |  | 0.500 |  | 0.500 |  | 0.373 |  | 0.627 |  | 0.594 |  | 0.003 |  | 0.034 |  | -0.072 |  | 0.096 |

*Note:* P(incl) = prior inclusion probability; P(excl) = prior exclusion probability; P(incl|data) = posterior inclusion probability; P(excl|data) = posterior exclusion probability; BF_inclusion_ = Bayes factor inclusion. The regression equation for the most parsimonious model is **parvalbumin proportion = 0.297 + (-0.21*diagnosis)** where a diagnosis of Down syndrome = 1 and control = 0.

**Supplementary table 20**. Descriptive statistics for parvalbumin Bayesian regression model

|  | | | | | | |
| --- | --- | --- | --- | --- | --- | --- |
|  | | **N** | | **Mean** | | **SD** |
| Parvalbumin proportion |  | 27 |  | 0.297 |  | 0.248 |
| Age |  | 27 |  | 52.074 |  | 13.023 |
| Aβ42 |  | 27 |  | 0.743 |  | 1.030 |

| **Supplementary table 21**. Calbindin proportion Bayesian regression model comparison. | | | | | | | | | | |
| --- | --- | --- | --- | --- | --- | --- | --- | --- | --- | --- |
| **Models** | | **P(M)** | | **P(M\|data)** | | **BF_M_** | | **BF_10_** | | **R²** |
| Null model |  | 0.250 |  | 0.005 |  | 0.015 |  | 1.000 |  | 0.000 |
| Diagnosis |  | 0.083 |  | 0.488 |  | 10.469 |  | 289.631 |  | 0.491 |
| Diagnosis + Age + Aβ42 |  | 0.250 |  | 0.192 |  | 0.713 |  | 38.026 |  | 0.512 |
| Diagnosis + Age |  | 0.083 |  | 0.144 |  | 1.845 |  | 85.308 |  | 0.497 |
| Diagnosis + Aβ42 |  | 0.083 |  | 0.139 |  | 1.782 |  | 82.824 |  | 0.495 |
| Age + Aβ42 |  | 0.083 |  | 0.024 |  | 0.274 |  | 14.433 |  | 0.394 |
| Aβ42 |  | 0.083 |  | 0.006 |  | 0.071 |  | 3.809 |  | 0.222 |
| Age |  | 0.083 |  | 0.001 |  | 0.016 |  | 0.880 |  | 0.093 |

*Note:* P(M) = prior model probability; P(M|data) = posterior model probability; BF_M_ = change from prior model odds to posterior model odds; BF_10_ = Bayes factor for each model compared to the null model.

**Supplementary table 22**. Parvalbumin proportion posterior summaries of coefficients

|  | | | | | | | | | | | | | | | | | | |
| --- | --- | --- | --- | --- | --- | --- | --- | --- | --- | --- | --- | --- | --- | --- | --- | --- | --- | --- |
|  | | | | | | | | | | | | | | | | **95% Credible Interval** | | |
| **Coefficient** | | **P(incl)** | | **P(excl)** | | **P(incl\|data)** | | **P(excl\|data)** | | **BF_inclusion_** | | **Mean** | | **SD** | | **Lower** | | **Upper** |
| Intercept |  | 1.000 |  | 0.000 |  | 1.000 |  | 0.000 |  | 1.000 |  | 0.313 |  | 0.038 |  | 0.244 |  | 0.404 |
| Diagnosis |  | 0.500 |  | 0.500 |  | 0.963 |  | 0.037 |  | 25.849 |  | -0.293 |  | 0.108 |  | -0.451 |  | 0.000 |
| Age |  | 0.500 |  | 0.500 |  | 0.361 |  | 0.639 |  | 0.566 |  | 9.365×10^-4^ |  | 0.003 |  | -0.002 |  | 0.008 |
| Aβ42 |  | 0.500 |  | 0.500 |  | 0.362 |  | 0.638 |  | 0.568 |  | -0.014 |  | 0.037 |  | -0.136 |  | 0.037 |

*Note:* P(incl) = prior inclusion probability; P(excl) = prior exclusion probability; P(incl|data) = posterior inclusion probability; P(excl|data) = posterior exclusion probability; BF_inclusion_ = Bayes factor inclusion. The regression equation for the most parsimonious model is **calbindin proportion = 0.313 + (-0.293*diagnosis)** where a diagnosis of Down syndrome = 1 and control = 0.

**Supplementary table 23**. Descriptive statistics for calbindin Bayesian regression mode

|  | | | | | | |
| --- | --- | --- | --- | --- | --- | --- |
|  | | **N** | | **Mean** | | **SD** |
| Calbindin proportion |  | 26 |  | 0.313 |  | 0.264 |
| Age |  | 26 |  | 51.385 |  | 12.769 |
| AB42 |  | 26 |  | 0.770 |  | 1.041 |
